# Unbiased Strain-Typing of Arbovirus Directly from Mosquitoes Using Nanopore Sequencing: A Field-forward Biosurveillance Protocol

**DOI:** 10.1101/183780

**Authors:** Joseph A. Russell, Brittany Campos, Jennifer Stone, Erik M. Blosser, Nathan Burkett-Cadena, Jonathan Jacobs

## Abstract

The future of infectious disease surveillance and outbreak response is trending towards smaller hand-held solutions for point-of-need pathogen detection.^1–4^ Although recent advances have paved the way for these technologies to include sequencing of pathogens directly from clinical samples, the ability to carry out unbiased sequencing for pathogen discovery and subtyping directly from environmental samples has yet to be demonstrated with hand-held platforms.^5^ Products such as the two3 qPCR system from Biomeme Inc., as well as the MinION from Oxford Nanopore Technologies, have generated renewed prospects for point-of-need diagnostics and near real-time environmental testing and characterization of viral and microbial pathogens. Here, samples of *Culex cedecei* mosquitoes collected in Southern Florida, USA were tested for Venezuelan Equine Encephalitis Virus (VEEV), a previously-weaponized arthropod-borne RNA-virus capable of causing acute and fatal encephalitis in animal and human hosts. A single 20-mosquito pool tested positive for VEEV by real-time reverse transcription quantitative PCR (RT-qPCR) on the Biomeme two3. The virus-positive sample was then subjected to unbiased metatranscriptome sequencing on the MinION and determined to contain Everglades Virus (EVEV), a strain of VEEV transmitted exclusively by *Culex cedecei* in South Florida. The result was confirmed on “gold standard” thermocyclers and sequencing machines, and comparison to nanopore results is discussed. Our results demonstrate, for the first time, the use of unbiased sequence-based detection and subtyping of a high-consequence biothreat pathogen directly from an environmental sample using field-forward hardware and protocols. The further development and validation of methods designed for field-based diagnostic metagenomics and pathogen discovery, such as those suitable for use in mobile “pocket laboratories”, will address a growing demand for public health teams to carry out their mission where it is most urgent: at the point-of-need.^6^

## INTRODUCTION

With increasing accessibility of metagenomics- and metatranscriptomics-based analyses (meta-omics), clinicians and researchers have begun to embrace the technology as a means of detection for unknown etiological agents of disease.^5,7–10^ In addition, metagenomics has an emerging role in environmental biosurveillance across multiple mission contexts including bioterrorism defense^11^, epidemiological public health^12–14^, water-quality monitoring^15^, and agriculture/food safety^16–18^. In comparison to PCR-based amplicon assays, metagenomics has the added value of not requiring *a priori* knowledge of a target (i.e., unbiased), delivers functional genomic information of constituent organisms in a sample (in addition to detection), and provides an estimate of their relative abundance. However, the benefits of this information are inextricably dependent on the quality of sample extraction and sequencing reads, the depth of sequencing and titer-level of the etiological agent, the comprehensiveness of reference databases, and the power and suitability of back-end computational equipment and bioinformatics analysis. Additionally, metagenomic sequencing on second-generation sequencing technology typically requires more than a 24 hour time investment on non-portable machines. Consequently, field-forward biosurveillance has been limited to primarily PCR-based assays^19–21^ or antibody hybridization technologies^22–25^, which have been the first molecular biology hardware to reach a portable, hand-held form factor.

Recent development in nanopore technology, pioneered by Oxford Nanopore Technologies, Inc. (ONT) with their MinION sequencing device, has opened the possibility of bringing the power of metagenomics to virtually any environment in the world. The MinION is a pocket-sized, USB-powered nanopore sequencing platform, weighing less than 100 grams, yet capable of up to 20 GB of ultra-long read (>100kb) sequence data.^26,27^ The device’s ultra-portability has been leveraged to perform in-field metagenomic characterization of environments ranging from the deep subsurface^28^ to the Antarctic Dry Valleys.^29^ But perhaps the most compelling application of the MinION platform is the improvement of pathogen surveillance and diagnostics, and subsequently, health outcomes, for the world’s most disadvantaged populations. The small footprint of the MinION, and other hand-held molecular biology hardware, is particularly important for austere settings with limited access to the critical infrastructure often required for traditional diagnostics and biosurveillance assays. Routine, point-of-sampling detection, phylogeny, and genomic characterization of microbial and viral pathogens from clinical *and* environmental samples stands to fundamentally change public health practices.^30–33^ Critically, nanopore sequencing has the added benefit of real-time analysis^34^, allowing sample-to-answer intervals that match clinically relevant timeframes. Recent work has demonstrated the efficacy of nanopore sequencing in RNA-based metatranscriptomic detection of viral pathogens from human blood samples.^35,36^ More recently, single-nucleotide polymorphism (SNP) detection was demonstrated on the MinION, using PCR amplicons of short tandem repeats, for the purposes of forensic genotyping.^37^ During the Zika Virus (ZIKV) outbreak of 2015-2016 in Brazil, several groups used nanopore sequencing of RT-qPCR amplicons from mosquito samples to track incidence of ZIKV infection and study ZIKV vector dynamics.^38,39^ And recently, an Australian group demonstrated the use of nanopore sequencing for whole genome sequencing of Ross River Virus, directly from a single mosquito under laboratory control conditions.^40^ However, to date, there have been no reports of unbiased (non-PCR) strain-level detection of specific organisms-of-interest directly from environmental sample matrices (e.g., non-clinical, non-sterile, nonlaboratory derived) using nanopore sequencing. This is likely due to the lower sequencing depth of nanopore data relative to second-generation sequencing machines, and subsequent detection of predominantly host genomic material. Over-coming this challenge will enable genome-based biosurveillance without the constraint of PCR primer design and optimization. This would be particularly useful for monitoring arbovirus and other viral hemorrhagic fever (VHF) vectors in hot-spot regions throughout the world where frequent epizootic events threaten the health of human populations. Often, the pathogens responsible for these events are RNA viruses with small genomes and high mutation rates, rendering the maintenance of high-fidelity primer sets an ongoing challenge.

An example of such a pathogen can be found in the Americas. Venezuelan Equine Encephalitis Virus (VEEV) is a positive-sense single-stranded RNA virus with an approximately 11.4 kilobase (KB) genome. An important human and equine pathogen that has previously been weaponized, VEEV is categorized as an overlap Select Agent by the U.S. government due to its pathogenicity to both humans and livestock. VEEV is responsible for the most persistent recurrent outbreaks of New World alphaviruses in the *Togaviridae* family^41^. In humans, VEEV causes a non-specific febrile illness, with onset of symptoms (fever/chills, malaise, tachycardia) after a 2 to 5-day incubation period. More severe cases (<1% in humans) will result in encephalitis, and eventually, death 5 to 10 days after infection^42^. It has been determined that some enzootic equine-avirulent VEEV strains can alter their serotype, and range of both mosquito vector and vertebrate host, through mutations in the genes encoding the E2 envelope glycoprotein^43^. Adaptation to equines results in extremely high viremia (>10^7^ PFU/ml), leading to a greater chance of human disease, and highlighting the role of genome-based strain tracking for public health purposes. An enzootic, sylvatic strain of VEEV (subtype II) circulates in and around the Everglades region of Southern Florida. Commonly known as Everglades virus (EVEV), this VEEV subtype is exclusively transmitted by the mosquito species *Culex (Melanoconion) cedecei*, with cotton rats and cotton mice as its primary vertebrate host.^41,44,45^ Surveys in the 1960’s and 1970’s indicated high seroprevalence of EVEV antibodies in humans residing in Southern Florida (>50% amongst Seminole Native Americans living north of Everglades National Park)^46–48^, and it has been suggested that EVEV may be an important, unrecognized cause of human illness in the region.^44^

In this study, we successfully demonstrate field-ready protocols for sample collection, RNA extraction, reverse transcription quantitative PCR amplification, eukaryote host genome depletion, and nanopore sequencing of a mosquito sample metatranscriptome for the purposes of arbovirus biosurveillance (Figure 1). We report the first use of nanopore sequencing to detect, and strain-type, an arbovirus directly from field-trapped mosquitoes using a metatranscriptome approach. The EVEV-positive sample was processed using current “gold standard” platforms (e.g., CFX-96, Illumina MiSeq) to benchmark differences in results with more conventional methods. This work demonstrates the practical utility of field-able, hand-held thermocyclers and nanopore sequencing devices for unbiased strain-level detection of high-titer arboviruses from complex, environmental sample matrices.

**Figure 1:**
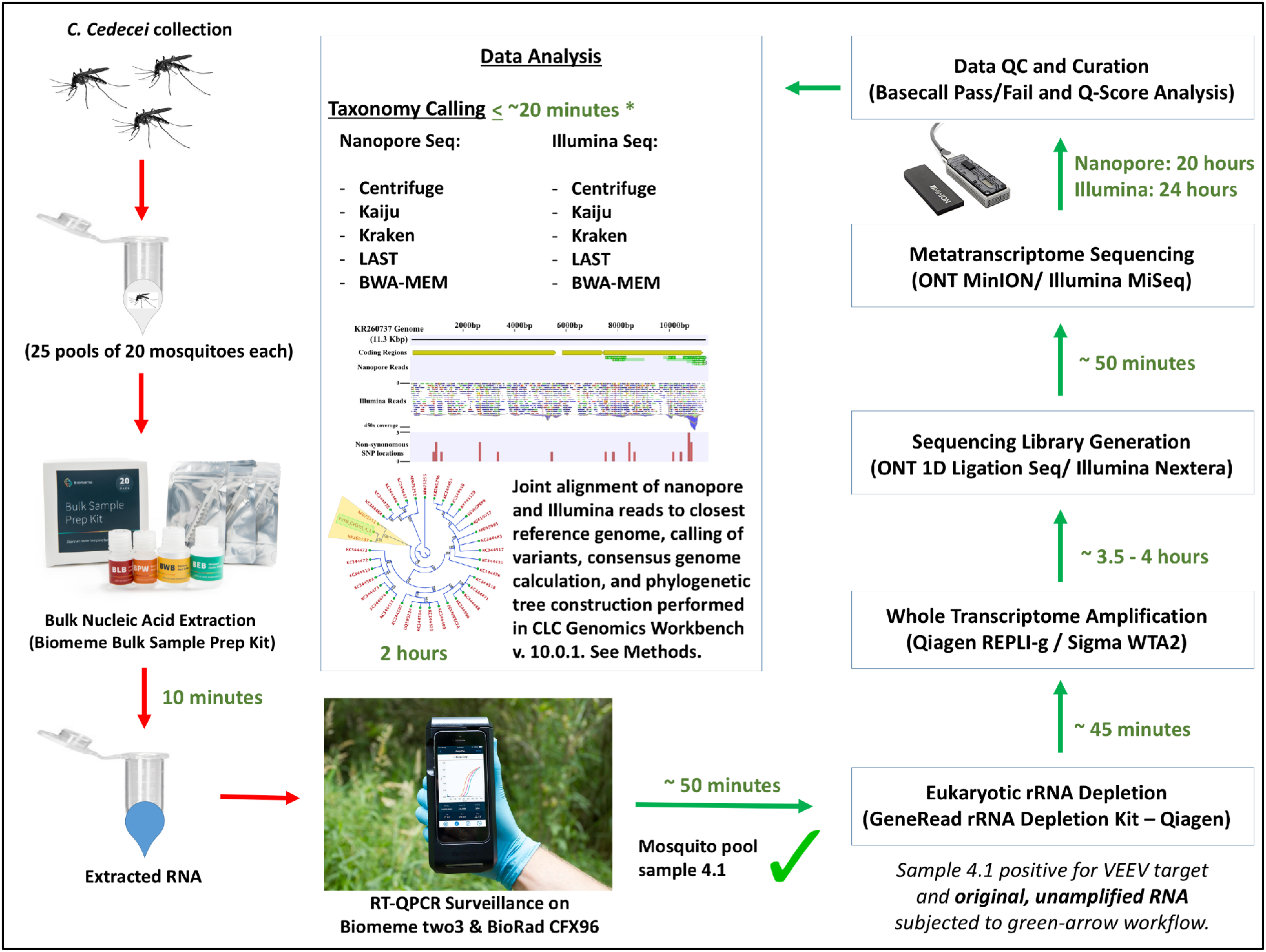
Overview of comparison between experimental field-based workflow and “gold-standard” methods. Times to perform individual steps are listed in green. Times listed are for a single hypothetical sample. Multiple samples can be processed at some steps with minimal impact on process time (e.g., rRNA depletion, WTA) while other steps will have more substantial increase in processing time with additional samples (e.g., nucleic acid extraction, data analysis). (*) indicates computational analysis time for nanopore data on a hyperthreaded quad-core, 32GB RAM computing system (Intel NUC Skull Canyon). Kraken could not be deployed on the Intel NUC due to memory limitations, and was run on a 16-core, 128GB RAM high-performance computing cluster.

## METHODS

### Sample Collection

Mosquito traps (CO_2_/light-baited) were set for overnight collection near carbonate dissolution pools in a forested environment, adjacent to Canal 111E in Homestead, FL, USA, on October 17^th^, 2016. The first clinical case of Everglades virus in humans was likely acquired while fishing along C-111 canal.^49^ The sampling site was located at 25.4078, -80.5237, approximately 3.8 miles from the Ingraham Highway entrance to Everglades National Park, FL, USA. Several thousand mosquitoes were collected. Female *Culex cedecei* individuals were visually sorted and separated via light microscopy inspection into their own sample pools of 20 individuals per 1.5 ml Eppendorf tubes. Twenty-five (25) sample pools were sorted, for a total of 500 female *Culex cedecei* mosquitoes.

### Sample Extraction

Bulk nucleic acids were extracted from each mosquito pool individually using the Bulk Nucleic Acids Field Extraction kit from Biomeme, Inc. (Philadelphia, PA, USA). The manufacturer’s protocol was followed, with slight modifications. Each 20-mosquito pool was mashed in 1.5 ml tube with kit-provided pestle for 1 minute. 50 μl of Biomeme Lysis Buffer (BLB) was added to the tube and mashing continued for an additional minute. 450 μl of BLB was added and mashing continued for an additional 30 seconds. The tube was then vortexed for 1 minute, then centrifuged for 1 minute at 5,000 × *g* to pellet course debris. Subsequently, 500 μl of supernatant was transferred to 1000 μl aliquot of BLB. The supernatant/BLB mix was briefly vortexed to mix. The Biomeme syringe extraction column was assembled and the entire supernatant/BLB mix was drawn up through the column, and then expelled slowly three times. Next, the entire volume of a 500 μl aliquot of Biomeme Protein Wash solution (BPW) was drawn up through the column and expelled slowly. Then, the entire volume of a 750 μl aliquot of Biomeme Wash Buffer (BWB) was drawn up through the column and expelled slowly. After expelling the BWB, the column was pumped continuously (air-dried) without any reagents until no buffer was spraying from the tip into the collection vial and there were minimal droplets in the column’s tubing. Finally, 200 μl of Biomeme Elution Buffer (BEB) was drawn into the column and allowed to incubate for 1 minute at room temperature. The BEB containing eluted total nucleic acids (TNA) was then expelled into a fresh 1.5 ml tube.

### RT-qPCR

Each mosquito pool RNA extract was queried with a quantitative real-time reverse transcription PCR assay specific for Venezuelan Equine Encephalitis Virus (VEEV), using the SuperScript^™^ III Platinum^®^ One-Step Quantitative RT-PCR System from Invitrogen (Waltham, MA, USA). The master mix contained, per 25 μl reaction; 5.25 μl dH_2_O, 12.5 μl 2X reaction mix, 0.5 μl RNaseOUT^™^ ribonuclease inhibitor, 0.5 μl Superscript III^™^ RT/Platinum^®^ Taq polymerase, 0.5 μl VEEV forward primer, 0.5 μl VEEV reverse primer, and 0.25 μl of VEEV taq-man probe. 5 μl sample RNA was added to each reaction. The RT-qPCR reactions for all 25 samples, plus a positive control and a no-template negative control, were run on both the Biomeme two3 handheld qPCR machine and the BioRad CFX96 Touch^™^ Real-time PCR detection system with the following cycling conditions; 50°C for 15 minutes, 95°C for 2 minutes, then 50 cycles of 95°C for 15 seconds and 60°C for 1 minute. A single mosquito pool (sample 4.1) was positive for VEEV on both the Biomeme two3 machine and the CFX96. Primer and probe sequences available upon request.

### Generation of Metatranscriptomes

We processed two samples for metatranscriptome sequencing; the single sample that tested positive for EVEV (4.1) and a sample that was negative for EVEV (1.1). Following the manufacturer’s protocol, the GeneRead rRNA Depletion Kit (Qiagen, Inc., Hilden, Germany) was used to reduce the burden of *C. cedecei* vector DNA and RNA. Depleted samples were then processed with the REPLI-g Single Cell Whole Transcriptome Amplification (WTA) kit (Qiagen, Inc.), according to the manufacturer’s protocol. After review of the REPLI-g nanopore data, it was determined that a comparison to another WTA method for nanopore metatranscriptome sequencing was prudent (additional details in Results). Following the manufacturer’s protocol, we also processed the raw TNA sample with the WTA2 Complete Whole Transcriptome Amplification kit from Sigma-Aldrich Inc. (St. Louis, MO, USA). WTA products (from either kit) were purified with Agencourt (Beverly, MA, USA) AMPure^®^ beads as follows; 1.8x the eluted WTA product volume (54 μl) of AMPure beads was added to the WTA-product (30 μl) and pipette-mixed 10 times. The reaction was placed on a magnetic stand for 10 minutes and the cleared solution was aspirated away. The cDNA-bound magnetic beads were washed 2x with 200 μl 70% ethanol and allowed to air dry for 5 minutes. 40 μl of dH_2_O was added to the washed beads and pipette-mixed 10 times. The sample was placed back on the magnetic stand for 10 minutes. The purified, eluted cDNA was transferred to a fresh 1.5 ml tube and quantified with a Qubit flourometer (ThermoFisher Sci., Waltham, MA, USA) according to the manufacturer’s protocol.

### Nanopore sequencing of Metatranscriptomes

The WTA products from virus-positive sample 4.1 and virus-negative sample 1.1 were prepared for nanopore sequencing using ONT’s 1D Ligation Sequencing library preparation kit (SQK-LSK108), following the manufacturer’s protocol. The library was loaded onto an R9.4 flow cell. Two separate flow cells were used for each sample for sequencing of the REPLI-g generated samples. For the Sigma WTA2 generated samples, the same flow cell was re-used for sample 1.1 (virus-negative sample) *after* sequencing sample 4.1 (virus-positive sample) and flushing/washing with the ONT Flowcell Wash Kit (EXP-WSH002). For all WTA products, the NC_48Hr_Sequencing_Run_FL0-MIN106_SQK-LSK108_plus_Basecaller.py script was used for collecting data. The REPLI-g 4.1 sample was run for approximately 26.5 hours, with a total of 1142 channels with active pores detected during the pre-run mux scan. The REPLI-g 1.1 sample was run for approximately 12 hours, with a total of 1465 channels with active pores detected during the pre-run mux scan. The Sigma WTA2 4.1 sample was run for approximately 20 hours, with a total of 1168 channels with active pores detected during the pre-run mix scan. The Sigma WTA2 1.1 sample was run for approximately 7.5 hours, with a total of 551 channels with active pores detected during the pre-run mux scan.

### Illumina sequencing of Metatranscriptomes

The WTA product from virus-positive sample 4.1 was prepared for Illumina sequencing using Illumina’s Nextera XT library prep kit (FC-131-1024), following the manufacturer’s protocol through the library clean-up step. Manual normalization was performed following DNA quantitation of the CAN product using the Qubit fluorometer to ensure a sufficient quantity of library was generated. The library was then diluted using a conversion factor of 2 to 2nM and pooled with other libraries. The libraries were added to a cartridge at a final loading concentration of 12pM using a MiSeq Reagent Kit V2 (MS-102-2002). A 2 × 151 paired-end run was performed on the Illumina MiSeq system (SY-410-1003) using the FASTQ only workflow.

### Bioinformatics and Data Analysis

#### Nanopore Data

Nanopore reads were basecalled using the local basecalling algorithm in MinKNOW version 1.4.3. FAST5 files of basecalled reads were converted to FASTA files using*poretools*.^50^ The FASTA files from each sequenced sample (4.1 and 1.1) were queried for EVEV/VEEV using several kmer-based metagenomics taxonomy callers, including Kraken^51^, Kaiju^52^, and Centrifuge.^53^ Two full-length read-mapping alignment tools, LAST^54^ and BWA-MEM^55^, were also tested. For computational resource and analysis time considerations, these read-mapping tools were deployed with a custom database of 144 VEEV genomes (rather than the larger RefSeq-sized databases of the kmer tools). See *Supplementary Material* for specific parameters called for each tool, as well as a list of accession numbers for all VEEV references in the custom database. The SAM alignment file generated via BWA-MEM mapping (with ‘-x ont2d’ flag called) to the custom VEEV database was imported into CLC-Genomics Workbench version 10.0. 1 (CLC) and converted to tracks for visualization purposes and exploratory analysis in comparison to Illumina MiSeq reads mapping to the same VEEV genomes.

#### Illumina Data

Sequencing reads from sample 4.1 were trimmed to a Q=30 quality score in CLC and analyzed for total taxonomic composition using *kraken, kaiju*, and *centrifuge* (See *Supplementary Material* for specific parameters for each tool). In addition, reads were mapped to the custom database of VEEV genomes with BWA-MEM and in CLC. Variants were called in CLC using the Basic Variant Detection Tool, which makes no assumptions about the underlying data. Sixteen (16) high frequency (HF) variants were called using this tool such that non-specific matches were ignored, a minimum of 30x coverage was required, and the variant called was required to be 100% penetrant (homozygous) with a minimum Q30 quality score. Three HF variants were predicted to result in amino acid changes. Low frequency (LF) variants, 60 in total including HF variants, were also called in a similar manner but had a 30x minimum coverage, with the variant allele being a minimum 10% frequency and 10x coverage at Q30 or above. A total of 18 LF variants were predicted to result in amino acid changes. A consensus genome was generated from the top two VEEV genomes recruiting the most reads, and the Illumina reads themselves. The consensus genome, called ‘EVG-2016_CxCdci_4_1’, was included in a multiple sequence alignment (MSA) with the database of 144 VEEV genomes. A pruned phylogenetic tree was constructed using the 30 closest relatives. The chosen tree construction was the neighbor-joining method^56^ and the nucleotide distance measure used was Jukes-Cantor.^57^ The tree was validated with 1,000 bootstrap replicates.

#### Data Availability

All raw sequence data from this work can be found at NCBI under BioProject PRJNA399278 (https://www.ncbi.nlm.nih.gov/bioproject/399278). Illumina data for Sample 4.1 is deposited with accession #XXXX. Nanopore data from REPLI-g amplified Sample 4.1 is deposited under accession # XXXX and Sample 1.1 is under accession # XXXX. Nanopore data from Sigma WTA2 amplified Sample 4.1 is deposited under accession # XXXX and Sample 1.1 is under accession # XXXX. *De-novo* assembled contigs for EVEV strain EVG-2016_CxCdci_4_1 were submitted to GenBank with accession # XXXX. (#XXXX = *Data currently being submitted to public archives*.)

## RESULTS

### RT-QPCR Arbovirus Surveillance

During our study, a single sample pool (Sample 4.1) tested positive for VEEV with a C_t_ value of 33.92 on the Biomeme two3 machine. The only sample that was positive for VEEV on the CFX96 system was also 4.1, with a C_t_ value of 30.63 (Figure 2). The Biomeme two3 device proved to be an effective, ultra-portable platform for initial triaging of mosquito samples in the field. While it could benefit from a higher throughput capacity, its small size and intuitive user interface render it a very capable field-forward molecular biosurveillance tool. It can also perform as a field-able heat-block and thermocycler for the steps in nanopore library generation that require such items.

**Figure 2:**
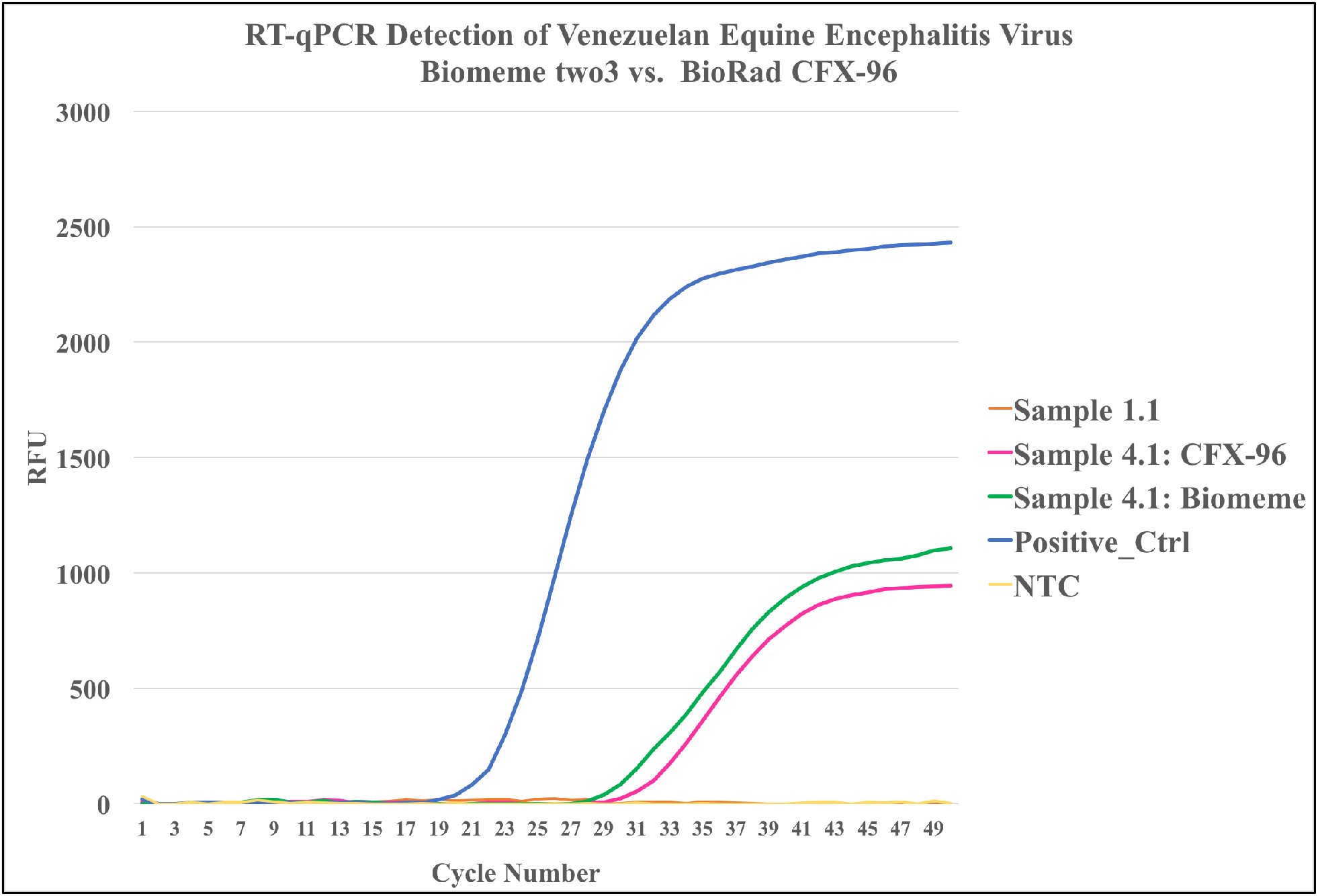
RT-qPCR results from mosquito pool sample 4.1 and 1.1. on the Biomeme two3 handheld thermocycler (green line) and the Bio-Rad CFX-96 benchtop thermocycler (pink line) in the laboratory. Sample 4.1 was positive for VEEV on both platforms; Biomeme C_t_ value = 33.92, CFX-96 C_t_ value = 30.63. Sample 1.1 was negative for VEEV on both platforms (CFX-96 data shown). NTC = No Template Control (nuclease-free H_2_O).

### Nanopore Sequencing (REPLI-g Single Cell WTA)

426,580 reads were successfully basecalled for the REPLI-g processed Sample 4.1; the average read length was 1,403 bp and the maximum read length was 21,258 bp. 106,040 reads were successfully basecalled for the REPLI-g processed Sample 1.1; the average read length was 2,038 bp and the maximum read length was 61,951 bp (Table 1). Detection of VEEV in 4.1 varied across several metagenomic taxonomy callers (Kraken, Kaiju, Centrifuge) that assign read-derived kmers to comprehensive genome databases (e.g., RefSeq). The two full-length read-mapping alignment tools (LAST, BWA-MEM) also varied in reported VEEV signal.

**Table 1:** Sequencing library information and VEEV/EVEV detection information across various analytical tools from both virus-positive (4.1) and virus-negative (1.1) samples. Numbers in the rows corresponding to taxonomic analysis tools indicate the number of VEEV/EVEV reads detected by that tool from the particular dataset. Asterisks (*) indicate analysis against a curated database of 144 VEEV and EVEV genomes, rather than the full RefSeq-sized database of the kmer tools (Kraken, Kaiju, Centrifuge). (GB) = gigabase

|  | 4.1 Illumina | 4.1 Nanopore (REPLI-g) | 1.1 Nanopore (REPLI-g) | 4.1 Nanopore (Sigma) | 1.1 Nanopore (Sigma) |
| --- | --- | --- | --- | --- | --- |
| # of sequencing reads | 6,429,832 | 426,580 | 106,040 | 212,192 | 71,355 |
| Avg. Read Length | 143 | 1,403 | 2,038 | 957 | 448 |
| Max. Read Length | 151 | 21,258 | 61,951 | 13,207 | 60,895 |
| No. of bases (GB) | 0.86 | 0.6 | 0.22 | 0.1 | 0.32 |
| Kraken | 796 | 1 | 0 | 2 | 0 |
| Kaiju | 2,420 | 10 | 0 | 17 | 1 |
| Centrifuge | 1,042 | 2 | 0 | 2 | 0 |
| LAST* | --- | 13 | 1 | 446 | 142 |
| BWA MEM, default* | 5,269 | 27 | 0 | 21 | 6 |
| BWA MEM, '-x ont2d'* | --- | 33 | 0 | 74 | 21 |
| CLC-Genomics* | 12,680 | --- | --- | --- | --- |

Kraken assigned a single nanopore read from sample 4.1 to VEEV, Centrifuge assigned 2 reads, and Kaiju assigned up to 10 reads. Kraken and Centrifuge offer less flexibility in parameter adjustment/loosening and were run with defaults as they were deemed acceptable for nanopore classification (i.e., Kraken’s *–min-hits* and Centrifuge’s *–min-hitlen* and –*min-totallen*). Kaiju allows greater flexibility in parameter adjustment. In our Kaiju submission script, we leveraged the ‘greedy mode’ and set the number of allowed mismatches to 10. We also lowered the minimum match score to 35 (from a default of 65). Running Kaiju with default parameters detected 9 VEEV reads in the 4.1 nanopore data. Loosening Kaiju’s parameters further than described above did not yield more than 10 VEEV reads. Of the direct read-mapping tools, BWA-MEM (with ‘-x ont2d’ flag passed) identified 33 VEEV reads from 4.1 nanopore data and LAST with nanopore-specific settings (see *Supplementary Material*) identified 13 VEEV reads. With default settings, BWA-MEM identified 27 VEEV reads in sample 4.1. A single read was classified as VEEV in virus-negative sample 1.1 by LAST, however, this was a secondary low-quality alignment (See *Supplementary Material*). The rest of the tools tested associated no reads with VEEV in sample 1.1 (Table 1).

For each read that was mapped to the VEEV database by BWA-MEM and LAST, the highest quality alignment was overwhelmingly one of two strains; EVG3-95 (KR260737) and Fe3-7c (AF075251). Both strains are the only Everglades Virus strains in the database of 144 VEEV genomes. The 33 reads aligned to VEEV by BWA-MEM were associated with 3 strains total; 19 reads to EVG3-95 (KR260737), 13 reads to Fe3-7c (AF075251), and 1 read to AG80-663 (AF075258, isolated in Argentina 1998). The 19 EVG3-95 reads covered 16% of the reference genome. The 13 Fe3-7c reads covered 10% of the reference genome (Table 2).

**Table 2:** Illumina and nanopore (REPLI-g vs. Sigma WTA2) read-mapping statistics of BWA-MEM with default settings (Illumina) and nanopore-specific settings (-x ont2d) for MinION data.

| **Illumina --- BWA-MEM (default)** |  |  |  |  |  |  |
| --- | --- | --- | --- | --- | --- | --- |
| VEEV Reference<br>Accession No. | Reference<br>length | Mapped<br>reads | Non-perfect<br>matches | % GC | Fraction of<br>reference covered | Total length<br>(zero coverage regions) |
| KR260737 | 11370 | 4073 | 2925 | 49.9 | 0.98 | 256 |
| AF075251 | 11395 | 1180 | 882 | 49.8 | 0.68 | 3594 |
| KC344490 | 11418 | 3 | 2 | 50.23 | 7.44E-03 | 11333 |
| KC344526 | 11418 | 3 | 1 | 50.22 | 9.63E-03 | 11308 |
| KC344471 | 11367 | 2 | 2 | 50.31 | 3.43E-03 | 11328 |
| KC344507 | 11135 | 2 | 1 | 50.18 | 7.63E-03 | 11050 |
| DQ390224 | 11412 | 1 | 0 | 50.19 | 4.73E-03 | 11358 |
| EEVCOMGEN | 11444 | 1 | 1 | 50.11 | 4.63E-03 | 11391 |
| EEVNSPEPB | 11446 | 1 | 1 | 49.76 | 7.16E-03 | 11364 |
| KC344475 | 11414 | 1 | 1 | 50.36 | 2.63E-03 | 11384 |
| KC344522 | 11412 | 1 | 1 | 50.10 | 4.38E-03 | 11362 |
| NC_001449 | 11444 | 1 | 1 | 50.11 | 4.63E-03 | 11391 |
| **Nanopore (REPLI-g) --- BWA-MEM (-x ont2d)** |  |  |  |  |  |  |
| VEEV Reference<br>Accession No. | Reference<br>length | Mapped<br>reads | Non-perfect<br>matches | % GC | Fraction of<br>reference covered | Total length<br>(zero coverage regions) |
| KR260737 | 11370 | 19 | 19 | 49.9 | 0.16 | 9502 |
| AF075251 | 11395 | 13 | 13 | 49.8 | 0.1 | 10263 |
| AF075258 | 11494 | 1 | 1 | 48.67 | 5.74E-03 | 11428 |
| **Nanopore (Sigma WTA2) --- BWA-MEM (-x ont2d)** |  |  |  |  |  |  |
| VEEV Reference<br>Accession No. | Reference<br>length | Mapped<br>reads | Non-perfect<br>matches | % GC | Fraction of<br>reference covered | Total length<br>(zero coverage regions) |
| KR260736 | 11495 | 45 | 45 | 49.73 | 0.02 | 11276 |
| KR260737 | 11370 | 25 | 25 | 49.9 | 0.55 | 5147 |
| AF075251 | 11395 | 4 | 4 | 49.8 | 0.11 | 10163 |

All REPLI-g nanopore reads mapping to Everglades Virus strain EVG3-95 via BWA-MEM aligned to the final ~4,000 basepairs of the 3’-region of the genome (Figure 3). This region encodes a sub-genomic 26S rRNA that is translated into a structural polyprotein which undergoes proteolytic cleaving to generate the viral capsid and the E2 and E1 envelope glycoproteins.^58^ A particularly high abundance of Illumina reads mapping to this region, and exclusive mapping of REPLI-g reads, is a likely indicator of an actively replicating viral infection since the 26S rRNA can only be transcribed from a full-length, negative sense RNA intermediate that itself can only be produced from the nsP1/nsP4 enzyme complex required for replication.^59^ RNA sequencing studies have recently shown that this region of the VEEV genome is also transcribed at significantly higher levels relative to the full length genomic RNA at the initial stages of infection. Thus, one can expect that a sample sequenced at this stage would have a high abundance of reads recruited to the 3’-region of the genome. This is observed in the alignment dynamics of both Illumina and REPLI-g nanopore reads. However, it should also be noted that the majority of VEEV-aligning REPLI-g generated nanopore reads were chimeric in nature. This can be visualized in the shade of green of REPLI-g nanopore reads aligning to Everglades Virus strain EVG3-95 (Figure 3). Darker green regions align to the reference, whereas lighter green regions do not.

**Figure 3:**
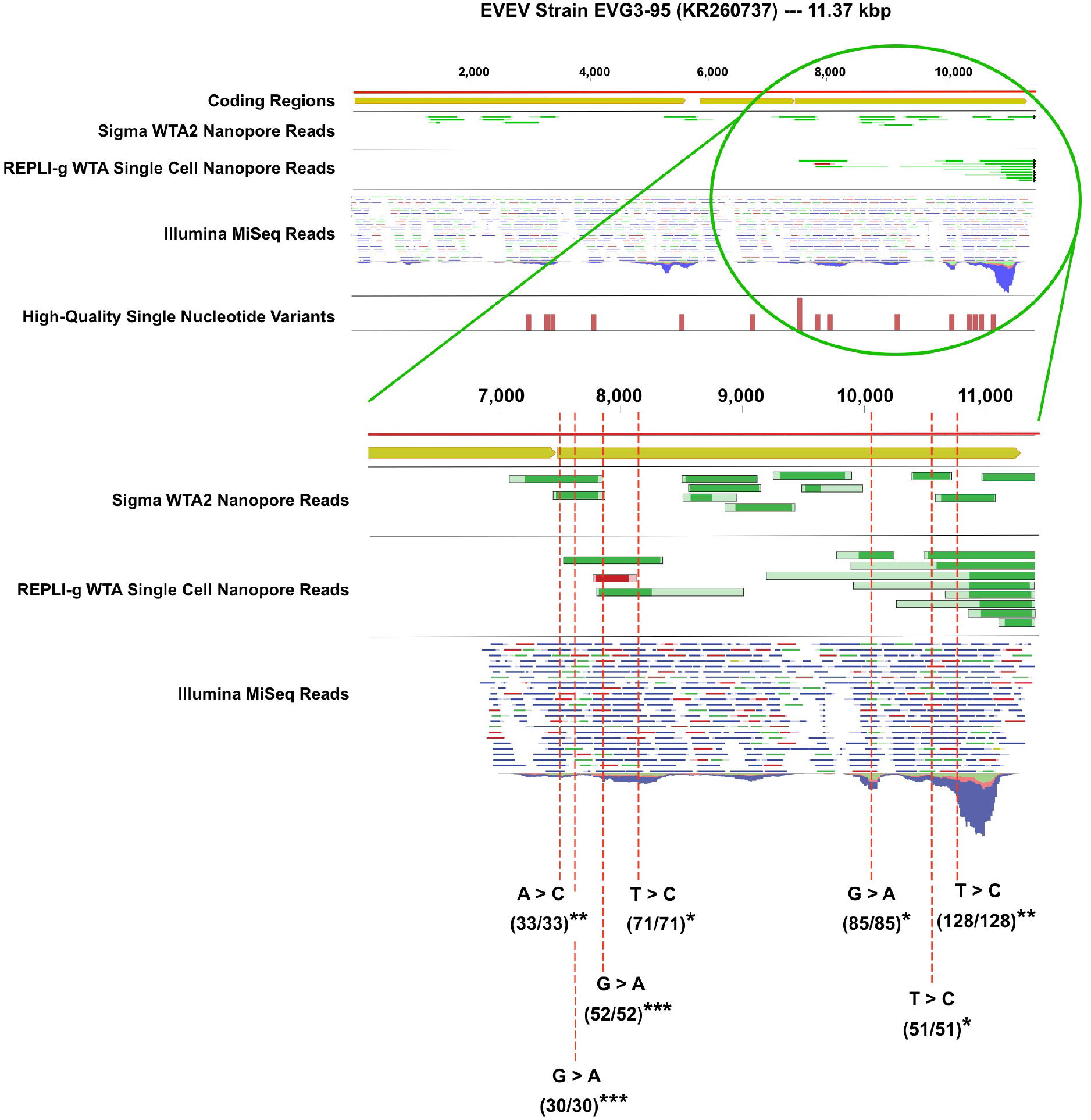
Mapping of Illumina reads and MinION nanopore reads generated through Sigma WTA2 and Qiagen REPLI-g to Everglades Virus strain EVG3-95. REPLI-g nanopore reads were exclusive to the final 4,000 basepairs (5’ → 3’), while Sigma nanopore reads were spread across the genome. Green shading of reads indicates alignment quality (dark regions align, light regions do not), and indicates ligated or otherwise generated chimeras. In the region where both sets of nanopore reads mapped, 7 out of 10 high-quality variants of 100% frequency detected by Illumina sequencing were also detected by nanopore sequencing. Only those variants detected by both Illumina and nanopore sequencing are shown. The ratio in parentheses below each variant is the ratio of Illumina reads containing the variant to Illumina read coverage at the specific location. The number of asterisks after the parentheses indicates how many nanopore reads also contained the same variant.

### REPLI-g Cell & Single Cell WTA Challenges for Nanopore Sequencing

Inspection of the alignments of REPLI-g generated nanopore reads that mapped to VEEV showed a high proportion of chimeric reads that only had a fraction of the read length aligning with any VEEV reference genomes. BLAST analysis of the remainder of the reads often hit to various mosquito species’ genomes (*Supplementary Figure 1A, 1B*), indicating combined vector/pathogen chimeric reads. Review of the specific chemistry of REPLI-g’s WTA process highlighted a step that is likely to be problematic for long-read sequencing technologies: namely, the ligation step. After complementary DNA (cDNA) is generated from RNA templates, the cDNA fragments are randomly ligated together to create longer molecules. This enhances the efficiency of the REPLI-g SensiPhi DNA polymerase during the multiple displacement amplification (MDA) reaction and, if used for populations of single cells from singular organisms, the impact on downstream quantification of transcripts is negligible. However, given the high efficiency and fidelity of the SensiPhi DNA polymerase, these kits have been attractive for *meta-omics* studies, with populations of multiple species, from low diversity, low biomass environments or investigations with minimal biological sample material^60–62^. For short-read sequencing technologies (i.e., Illumina), the confounding effects of the ligation step, and subsequent chimeric cDNAs, are negligible, or an acceptable trade-off for the efficacy of SensiPhi DNA polymerase. This is due to the low fraction of short reads that will, by chance, span a chimeric junction. With *long read* sequencing technologies, such as MinION or PacBio, this fraction is more likely to pose a challenge as long reads have a higher likelihood of spanning chimeric junctions. This is not necessarily problematic, depending on the use-case. For example, if the goal is simply detection of organisms of interest from a particular sample matrix in a biosurveillance context, then bioinformatics precautions can be set such that any existing signal will be recovered, despite the ligated fragments (e.g., reducing required fraction of reads that must align, reducing seed lengths, etc.). Indeed, informative recovery of Everglades Virus reads was observed with REPLI-g generated nanopore data (Table 1, Figure 3). However, in other analyses requiring high quality alignments (e.g., epidemiological strain mapping, genome finishing), these chimeras will present more of a problem. Ideally, no analyses are precluded from the generated data, so we selected another WTA kit to test that does not include the random ligations of cDNA fragments inherent to REPLI-g. We chose the WTA2 kit from Sigma-Aldrich.

### Nanopore Sequencing (Sigma WTA2)

212,192 reads were successfully basecalled for the Sigma WTA2 processed Sample 4.1; the average read length was 957 bp and the maximum read length was 13,207 bp. 71,355 reads were successfully basecalled for the REPLI-g processed Sample 1.1; the average read length was 448 bp and the maximum read length was 60,895 bp (Table 1). The same taxonomy-calling tools tested on the REPLI-g nanopore data were also tested on the Sigma WTA2 nanopore data, using the same settings. In Sample 4.1; Kraken detected 2 VEEV reads, Kaiju detected 17 VEEV reads, Centrifuge detected 2 VEEV reads, LAST detected 446 VEEV reads, and BWA-MEM detected 21 VEEV reads with default settings and 75 with the ‘-x ont2d’ flag passed. In Sample 1.1; Kraken detected 0 VEEV reads, Kaiju detected 1 VEEV read, Centrifuge detected 0 VEEV reads, LAST detected 142 VEEV reads, and BWA-MEM detected 6 VEEV reads with default settings and 21 with the ‘-x ont2d’ flag passed (Table 1). It is presumed that the higher incidence in VEEV reads detected in Sample 1.1 from Sigma WTA2 generated nanopore data is due to carry-over from insufficient washing and re-use of the flow cell after sequencing Sample 4.1. During REPLI-g testing, a fresh flow cell was used for each sample.

In contrast to the REPLI-g generated nanopore reads, Sigma WTA2 reads aligned to all coding regions of the EVG3-95 genome. Additionally, Sigma WTA2 generated nanopore reads showed lower rates of chimeric reads (Figure 3). This may be primarily due to the lack of the ligation step in the Sigma WTA2 protocol. However, various other key differences between the kits (e.g., SensiPhi vs. WTA2 polymerase activity, sequence composition of universal primers, etc.) are likely to contribute to observed differences in alignment dynamics for VEEV-associated reads. The 74 Sigma reads aligned to VEEV by BWA-MEM were associated with 3 strains; 45 reads to KR260736 (VEEV strain COAN5506, a 1967 equine isolate from Colombia), 25 reads to KR260737, and 4 reads to AF075251. While the numerical majority of Sigma WTA2 reads aligned to the Colombian strain, the coverage of this genome was much lower (2%) than that of the Everglades Virus strains (KR260737 - 55%, AF075251 - 11%) (Table 2). The total length of strain COAN5506 with zero read coverage is 11,276 basepairs out of a 11,495 bp genome, indicating stacking of approximately 219 bp reads at one location (Table 2).

### Illumina Sequencing

6,429,832 paired-end (2 × 151 bp) Illumina MiSeq reads were generated for Sample 4.1 and the average quality-trimmed read length was 143 bp (Table 1). The taxonomy-calling tools used on the nanopore datasets were also used on the Illumina data, however, default settings of each tool were used rather than nanopore-specific parameters (See Methods). Kraken identified 796 VEEV reads, Kaiju identified 2,420, Centrifuge identified 1,042, default BWA-MEM identified 5,269, and CLC identified 12,680 VEEV reads under default settings (Table 1). The highest represented VEEV strain in the Illumina data were the two Everglades virus (EVEV) strains; EVG3-95 (KR260737), followed by Fe3-7c (AF075251). These two strains accounted for 99.7% of all VEEV-associated reads as mapped by BWA-MEM (Table 2).

We observed a notable increase of Illumina reads mapping to the 26S sub-genomic RNA region, in the final 4.0 kb of the EVG3-95 genome, as was observed in the REPLI-g generated nanopore reads (Figure 3). While we predicted an active viral infection based solely on a limited number of nanopore reads, the higher density of Illumina reads mapping to the 26S region provides evidence of active EVEV replication in the 4.1 mosquito pool sample, rather than a latent infection or trace detection.

### Variant Detection in Nanopore and Illumina Data

From Illumina sequencing data, we observed 16 high-quality single nucleotide variants (SNVs) across the strain EVG3-95 genome (Table 3, Figure 3). 10 of these variants (~ 62%) were also detected in a nanopore sequencing read, regardless of which WTA-method was used. 10 SNVs were located in the 26S sub-genomic RNA region. Of these 10 variants, 7 were detected in a MinION nanopore read, 6 of the 7 were detected by a REPLI-g generated MinION read, and 3 of the 7 were detected by a Sigma WTA2 generated MinION read. Of the 6 SNVs in the first ~7 kb of the reference genome, 3 were *only* detected by Sigma WTA2 MinION reads. This data highlights the potential of the MinION nanopore sequencer to be leveraged for real-time, *unbiased*, SNV-level strain-tracking of arbovirus targets, directly from complex environmental samples.

**Table 3:**
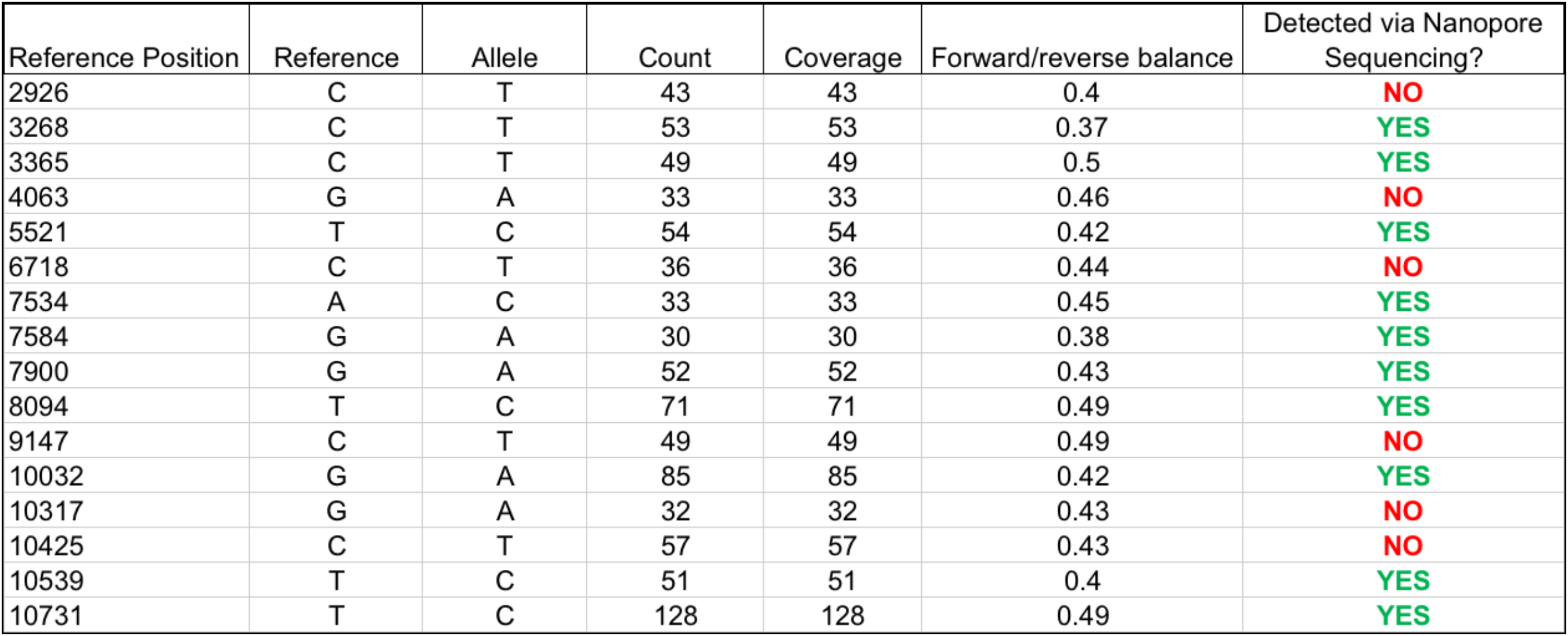
Metrics of high-quality (read count with variant / read coverage = 1 at given reference position) single nucleotide variants (SNVs) detected across the Everglades Virus strain EVG3-95 (KR260737) genome via Illumina MiSeq sequencing of virus-positive mosquito pool sample 4.1. 10 out of 16 SNVs (~62%) were detected via MinION reads.

### Phylogeny of EVEV-2016_CxCdci_4_1

Everglades virus strains belong to the Type-2 VEEV serogroup; a distinct phylogenetic group within the VEEV serocomplex. A consensus genome of the suspected strain of EVEV present in sample 4.1 was generated from the EVEV strain EVG3-95 genome, EVEV strain Fe3-7c genome, and the 12,680 Illumina reads mapping to these genomes in CLC. We label this consensus genome scaffold ‘EVG-2016_CxCdci_4_1’ (Figure 4). This name denotes the strain’s detection just outside of Everglades National Park in the autumn of 2016, the vector mosquito species (*Culex cedecei*), and the mosquito pool sample from this study that contained the virus (4.1). Phylogenetic analysis using full-length genomes of the 144 VEEV strains in our reference database clustered the EVG-2016_CxCdci_4_1 strain’s genome scaffold distinctly with the other EVEV strains and it appears more closely related to the EVG3-95 strain (KR260737) than the Fe3-7c strain (AF075251) (Figure 4).

**Figure 4:**
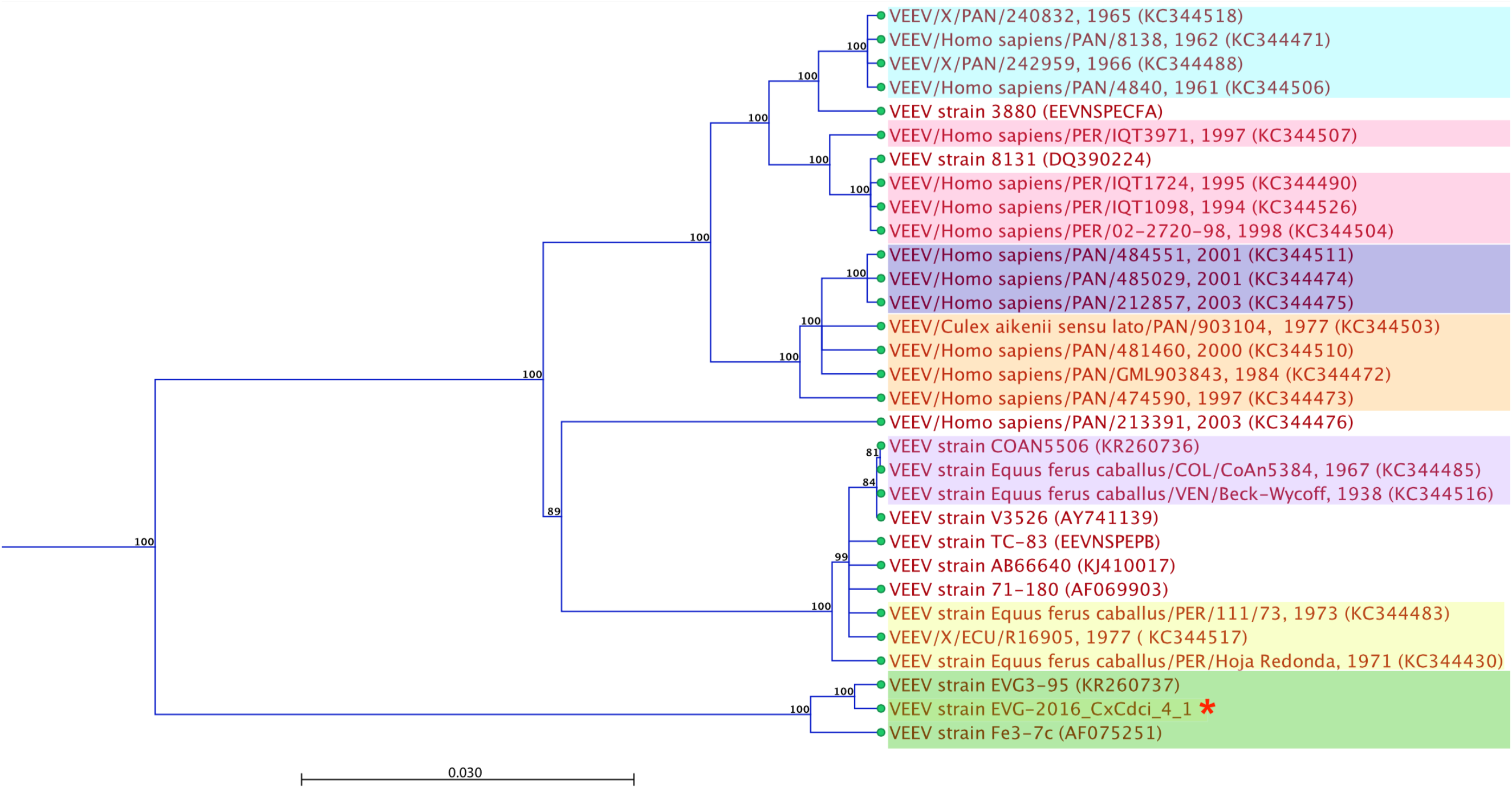
Whole genome phylogenetic tree of VEEV and EVEV strains and sub-types. The tree was clustered with the neighbor-joining method, invoking the Jukes-Cantor nucleotide substitution model, as implemented by CLC-Genomics Workbench version 10.0. 1. Bootstrap values >85 are shown (1,000 replicates). Scale bar indicates 3% nucleotide sequence divergence. Strains isolated from past epizootic events are indicated by colored boxes — Light blue (1961-1966, Panama), Pink (1994-1998, Peru), Dark blue (2001-2003, Panama), Orange (1977, 1984, 1997, 2000, Panama), Light Purple (1938-1967, Colombia/Venezuela), Yellow (1971-1977, Peru/Ecuador). Everglades Virus strains are highlighted in green. The red asterisk (*) indicates the consensus genome derived from KR260737, AF075251, and the Illumina sequencing reads generated in this study, denoted ‘EVG-2016_CxCdci_4_1’.

## DISCUSSION

Unbiased meta-omics approaches offer the ability to monitor the presence of nearly all potential pathogens in a single test. In geographic regions where several distinct pathogens can cause nearly identical febrile illness symptoms, the elimination of the need for multiple individual tests translates to reduced time for appropriate clinical or public health decisions to be made. Often, these same regions have limited capacity and infrastructure requirements to fully support brick-and-mortar laboratories and second-generation sequencing machines. Consequently, the prospect of unbiased meta-omics pathogen surveillance on devices as portable and low-maintenance as the MinION is a critical advantage that stands to fundamentally change the fight against emerging infectious diseases worldwide. However, challenges to the full realization of this potential remain.

When using unbiased meta-omics techniques, signal from the organism-of-interest is generally a small fraction of the total data output. Indeed, over 99% of nanopore reads from both sample 4.1 and 1.1 were annotated as ‘unclassified’ with Kaiju and Centrifuge analysis (*Supplementary File 1, Supplementary File 2*). The reference databases used by the metagenomics classifiers used are focused on microbial and viral species, so this result indicates that over 99% of the nanopore signal was (not surprisingly) from the eukaryote host (*Culex cedecei*). This was confirmed through BLAST analysis of several of the longest reads (*Supplementary Figure 1A, 1B*, and data not shown). The use of the GeneRead rRNA Depletion Kit was critical in this context, enabling sufficient host depletion for detection of EVEV RNA.

EVEV was detected in sample 4.1 using kmer-based taxonomy callers that leveraged RefSeq-sized databases, however, a more robust signal was observed when using read-mapping tools with target-specific databases (Table 1). While the use of targeted databases may preclude the reporting of other organisms that the MinION reads may map equally well to, it should not falsely inflate the presence of the organism-of-interest since read-mapping settings are fixed and each read is given an equal chance at mapping to each reference genome. Thus, we should expect the same number of reads associated with the organism-of-interest whether we are using all of RefSeq or a streamlined, targeted database. The key advantage in using streamlined databases targeting specific organisms-of-interest is that it enables read-mapping tools to be deployed on portable commodity computing systems (i.e., Intel NUC, Macbook Pro, etc.), further supporting the field-forward position of these types of analytical approaches. Importantly, field-forward researchers do not need to “choose” one or the other. Kmer-tools with comprehensive databases (i.e., Kaiju, Centrifuge) and read-mapping tools that query streamlined, targeted databases (i.e., BWA-MEM, LAST) can both be utilized effectively on portable computing systems. Therefore, there’s an advantage to installing Kaiju or Centrifuge, *and* BWA-MEM or LAST, onto any computing system meant for agnostic nanopore sequencing in the field, and using them in tandem with appropriate corresponding databases to conduct surveillance of broad groups of organisms.

Total analysis time for EVEV-positive sample 4.1, from sample collection through data analysis, was approximately 26 hours with MinION sequencing of the Sigma WTA2 product, compared to more than 30 hours with sequencing on the Illumina MiSeq (Figure 1). BWA-MEM, Centrifuge, Kaiju, and LAST were tested on a compact, portable commodity computing system (hyperthreaded quad-core, 32GB RAM Intel NUC Skull Canyon) running the Ubuntu 16.04 LTS operating system. These tools had rapid processing times for the nanopore data (≤~20 mins). The time-to-result for nanopore sequencing could have been reduced to between 3 and 6 hours if the original VEEV amplicon had been used as the sequencing material, and an internet connection was available for real-time taxonomy calling^34^. However, an agnostic approach demands extra time investment not required of amplicon sequencing due to the increased sequencing depth required to detect ultra-low abundance signals. We did not monitor the nanopore data in real-time, so it is not possible to determine exactly how long it took to detect an EVEV signal. Nonetheless, we were able to generate actionable biosurveillance data within a time frame amenable to enacting rapid response measures from public health entities (~1 day).

One aspect of ONT’s workflow that is critical for effective field deployment is their flow cell wash procedure. Minimizing the required amount of consumables that must be carried to remote field sites is still one of the primary challenges for mobilized deployment of nanopore sequencing, and so, the washing/flushing and re-use of the flow cells is an important feature of the MinION platform. In addition, when attempting to distinguish problematic or infectious samples from benign samples using agnostic sequencing, inter-run cross-contamination will be a confounding issue. We washed and re-used a R9.4 flowcell to sequence EVEV-negative, Sigma WTA2-processed sample 1.1 *after* we sequenced EVEV-positive, Sigma processed sample 4.1. We found low-level cross contamination of EVEV reads in sample 1.1 that we suspect originated from sample 4.1, despite following the ONT wash kit protocol exactly (Table 1). It is not suspected that this was trace signal of EVEV in sample 1.1 that was only detected with Sigma WTA2 processing since sample 1.1 was negative for VEEV/EVEV in the RT-qPCR assay (Figure 2). When a fresh R9.4 flowcell was used for each REPLI-g processed sample, no cross contamination was widely reported across tested taxonomy classification tools.

A list specific items (hardware, software, reagents, consumables, etc.) used to complete the work discussed here is given in the *Supplementary Material*. Taken together, these items can fit within a single, medium sized (40L) expidition-style backpack. This has not been lost on the research community and efforts to push nanopore-based molecular biosurveillance as far afield as possible have been prodigous^28,29,38,63,64^, including Low-Earth orbit^65^ and beyond.^66,67^ However, while carrying the items that are physically handled during sample processing is trivial, transporting the accompanying power and cold-chain logistical equipment has been more challenging and likely a primary factor preventing wider adoption of the technology in austere public health settings. The incredibly small footprint of the MinION is not fully empowered when one must also transport diesel generators, fuel, and mini-freezers as well. Development of intuitively designed, logistics-integrated, single-person portable laboratories will facilitate the future that the MinION’s form-factor inspires.

Future work will determine whether agnostic nanopore sequencing will be an effective biosurveillance tool on lower-titer pathogens - such as contaminated food samples or blood samples taken from sentinel wildlife populations. Our work likely benefited from the high-titer characteristic of VEEV infections and an actively replicating virus. However, the chemistry of ONT’s MinION flowcells and library preparation reagents remain under active development and improvements in both data yield and sequencing read quality are being released regularly^63^. We expect this to translate to unbiased strain-level detection of a wider array of organisms from even more challenging samples in the near future.

## CONCLUSIONS

Previous unbiased, meta-omics nanopore sequencing approaches to strain-specific target classification and SNV-calling have been limited to human blood, unknown isolates, or mixed culture sample matrices.^35,64^ In this study, we’ve pushed this capability to include complex biological sample matrices collected in the field - namely crushed mosquito pools collected from field traps. We describe a protocol that leverages ultra-compact hardware (e.g. the Biomeme two3, Intel NUC, and ONT MinION) to enable field-forward use of unbiased nanopore sequencing for the purposes of arbovirus biosurveillance. This work demonstrates the utility of nanopore sequencing for a wide array of public health and basic research use-cases in environmental biosurveillance. It is our hope that this work will further encourage the adoption of field-forward sequencing and bioinformatics to routinely bring the laboratory to the sample.

## Supplementary Material

### Taxonomy-caller parameters

~~~
------LAST-----
$ lastdb -Q 0 VEEV_reference_genomes VEEV_reference_genomes.fasta
$ lastal -s 2 -T 0 -a 1 -Q 1 -f BlastTab VEEV_reference_genomes data.fastq > data_alns.maf
$ grep “^[^#;]” data_alns.maf | awk -F ‘\t’ ‘{print $1}’ | sort | uniq -c | sort -nr | wc -l
~~~

~~~
------BWA-MEM------
$ bwa index VEEV_reference_genomes.fasta
$ bwa mem -x ont2d VEEV_reference_genomes.fasta data.fastq > data.sam
$ samtools view -bS data.sam > data.bam
$ samtools sort data.bam -o data_sorted.bam
$ samtools view -F 260 data_sorted.bam | cut -f 3 | sort | uniq -c | awk ‘{printf(“%s\t%s\n”, $2, $1)}’ > counts.txt
~~~

~~~
------CENTRIFUGE------
set -xeu
/src/centrifuge/centrifuge -x /src/centrifuge/indices/phv -U data.fastq -S data.out -p 16 --met-
stderr
~~~

~~~
------KRAKEN------
/home/bin/kraken \
      --preload \
      --db /home/src/kraken/full \
      --fastq-input \
      --threads 16 \
      --classified-out ./$DATA-class.fa \
      --unclassified-out ./$DATA-unclass.fa \
      --output ./$DATA-krakenout.txt \
      data.fastq
kraken-report --db /home/src/kraken/full ./$DATA-krakenout.txt > ./$DATA-kreport.txt
kraken-mpa-report --db /home/src/kraken/full ./$DATA-krakenout.txt > ./$DATA-mpkraken.txt;
~~~

~~~
------KAIJU------
set -xeu
~~~

~~~
kaiju -t /home/src/kaiju/bin/kaijudb/nodes.dmp -f /home/src/kaiju/bin/kaijudb/kaiju_db.fmi -i data.fastq -o ./data.kaiju.out -v -z 16 -a greedy -e 10 -s 35
~~~

~~~
addTaxonNames -t /home/src/kaiju/bin/kaijudb/nodes.dmp -n /home/src/kaiju/bin/kaijudb/names.dmp -i ./data.kaiju.out -o ./data.kaiju-names.out
~~~

~~~
kaijuReport -t /home/src/kaiju/bin/kaijudb/nodes.dmp -n /home/src/kaiju/bin/kaijudb/names.dmp -i ./data.kaiju.out -r species -o ./data.kaiju-names.out.summary
~~~

**Supplementary Figure 1:**
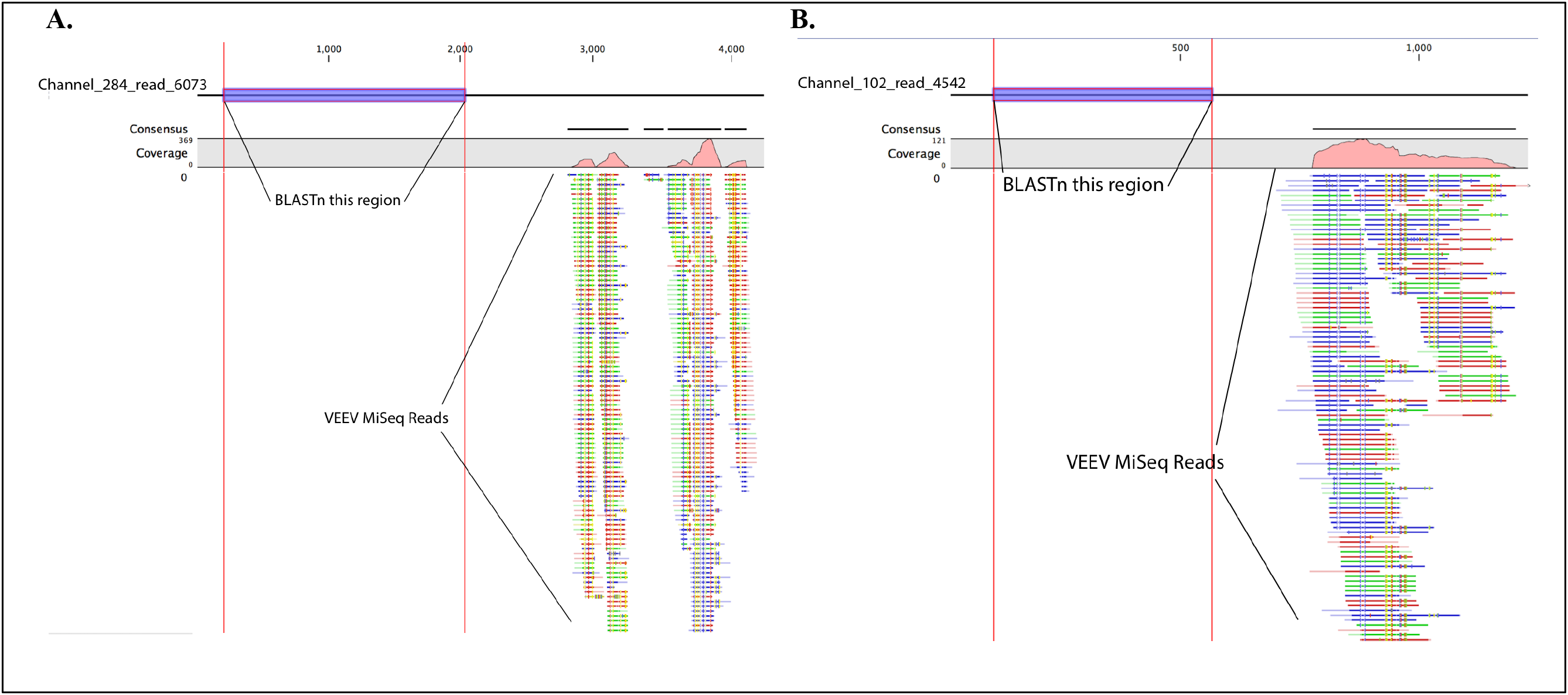
Two examples of chimeric REPLI-g generated nanopore reads from virus-positive mosquito pool sample 4.1. Illumina MiSeq reads that mapped to any strain in the custom VEEV database were isolated and re-mapped to REPLI-g generated nanopore reads that also aligned to VEEV references in CLC-Bio Genomics Workbench v. 10.0.1. One would expect generally uniform distribution of re-mapped MiSeq reads across VEEV-associated nanopore reads. However, MiSeq reads are observed to align with specific regions of nanopore reads, and are absent from other regions, indicating chimerism in REPLI-g generated nanopore reads. (A) The purple-highlighted region of Channel_284_read_6073 was BLASTed against the nt database and the highest associated hits were for the mosquito *Culex qainqaefasciatas* and *Drosophila* spp. (B) The purple highlighted region of Channel_102_read_4542 returned *C.ulex quinquefasciatus* and *Aedes aegypti* as top hits in its BLAST result.

#### List of Suggested Hardware and Consumables for Field-Forward Agnostic Nanopore Sequencing for the Purposes of Environmental Biosurveillance –––

1. *the Biomeme Bulk Nucleic Acid Extraction kit provides individually wrapped packets containing all plastic-ware needed for each sample (syringe assembly, tubing, plastic pestle, 2 ml elution tube) and 4 reagent bottles (2 × 15 ml, 2 × 30 ml)*
2. *the Biomeme two3 thermocycler (or other portable thermocycler: e.g., MIC, miniPCR, etc.) and accompanying 0.2 ml RT-qPCR tubes with lyophilized assays*
3. *the MinION nanopore sequencing device*
4. *required number of R9.4 flowcells*
5. *ONT sequencing library preparation kit (SQK-LSK108) and flow cell wash kit (EXP-WSH002), and associated reagents*
6. *GeneRead rRNA Depletion Kit*
7. *Sigma WTA2 Whole Transcriptome Amplification Kit*
8. *50 ml conical tube filled with 70% EtOH (1). 50 ml conical tube filled with 100% EtOH (1). 50 ml conical tube of molecular grade H_2_0 (1)*.
9. *mini centrifuge*
10. *P1000, P200, P20, P10 pipetman and tip boxes*
11. *magnetic tube stand*
12. *AMPure and Streptavidin beads (2 × 5 ml bottles)*
13. *Intel NUC Skull Canyon (32GB RAM, up to 2TB SSD, hyper-threaded quad-core, Ubuntu 16.04 LTS) and Bluetooth monitor/keyboard/trackpad*

#### List of VEEV Reference Genome Accession Numbers from Targeted Read-Mapping Database –––

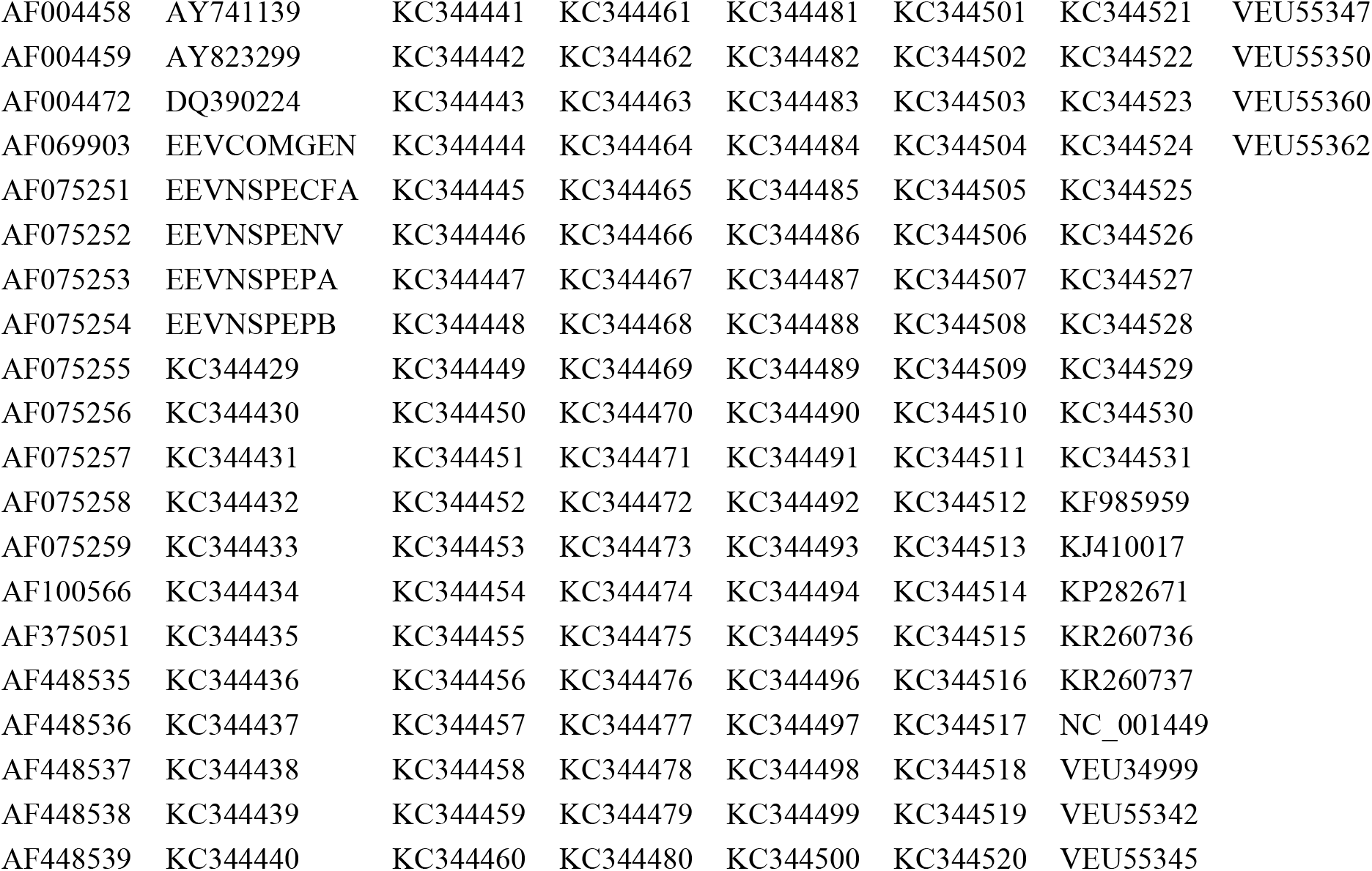

